# Advancing microRNA Target Site Prediction with Transformer and Base-Pairing Patterns

**DOI:** 10.1101/2024.05.05.592612

**Authors:** Yue Bi, Fuyi Li, Cong Wang, Tong Pan, Chen Davidovich, Geoffrey I. Webb, Jiangning Song

**Author notes:** Corresponding authors: Jiangning Song,; Fuyi Li,; Geoff Webb.

## Abstract

Micro RNAs (miRNAs) are short non-coding RNAs involved in various cellular processes, playing a crucial role in gene regulation. Identifying miRNA targets remains a central challenge and is pivotal for elucidating the complex gene regulatory networks. Traditional computational approaches have predominantly focused on identifying miRNA targets through perfect Watson-Crick base pairings within the seed region, referred to as canonical sites. However, emerging evidence suggests that perfect seed matches are not a prerequisite for miRNA-mediated regulation, underscoring the importance of also recognizing imperfect, or non-canonical, sites. To address this challenge, we propose Mimosa, a new computational approach that employs the Transformer framework to enhance the prediction of miRNA targets. Mimosa distinguishes itself by integrating contextual, positional, and base-pairing information to capture in-depth attributes, thereby improving its predictive capabilities. Its unique ability to identify non-canonical base-pairing patterns makes Mimosa a standout model, reducing the reliance on pre-selecting candidate targets. Mimosa achieves superior performance in gene-level predictions and also shows impressive performance in site-level predictions across various non-human species through extensive benchmarking tests. To facilitate research efforts in miRNA targeting, we have developed an easy-to-use web server for comprehensive end-to-end predictions, which is publicly available at http://monash.bioweb.cloud.edu.au/Mimosa/.

## 1. Introduction

Micro RNAs (miRNAs) are short non-coding RNAs, approximately 23 nucleotides long, critical in regulating gene expression at the post-transcriptional level (1,2). As part of the miRNA-induced silencing complex (miRISC), miRNAs form associations with Argonaute (AGO) proteins, facilitating their binding to target mRNA and leading to destabilization or translational repression (3). The specificity of miRNA targeting is primarily attributed to consecutive Watson-Crick (WC) base pairing between the miRNA seed region (positions 2-8) and complementary sites within the 3’ untranslated regions (3’ UTRs) of target mRNAs (4,5). The seed region contains well-known ‘canonical sites’, including 8mer, 7mer-m8, 7mer-A1, and 6mer, characterized by perfect Watson-Crick pairings (6). However, recent advances, particularly in high-throughput sequencing techniques like CLIP-seq, suggest that perfect seed matches are neither necessary nor sufficient for the functionality of the miRNA-target identification (7-9). It has been observed that imperfect base pairing also leads to a decrease in mRNA levels (10,11). These instances are typically referred to as ‘non-canonical sites’, as depicted in **Figure 1A**. For instance, functional interactions can tolerate a limited number of wobble pairs, bulges, and mismatches within the seed region (12,13). ‘Centered sites’ are a class of contiguous base-pairing of 11-12 nucleotides that starts at the 3^rd^ or 4^th^ nucleotide and extends to the central region of the miRNA (14). In addition, wobble or mismatch in the seed can be compensated by pairing with the 3’ region (positions 13-16) (15,16). Even perfect seed pairing may also be enhanced by complementary interactions within the 3’ region (17). Despite the discovery of these seed base pairings, fully unravelling the functional mechanism of miRNAs remains a challenge. One of the critical issues is to decipher the complexity of the miRNA regulatory network because one miRNA can target multiple different mRNAs and one mRNA can be regulated by many different miRNAs (18). Therefore, determining which miRNAs interact with which mRNAs, i.e., identifying miRNA targets, continues to be an important focus of current research.

**Figure 1.**
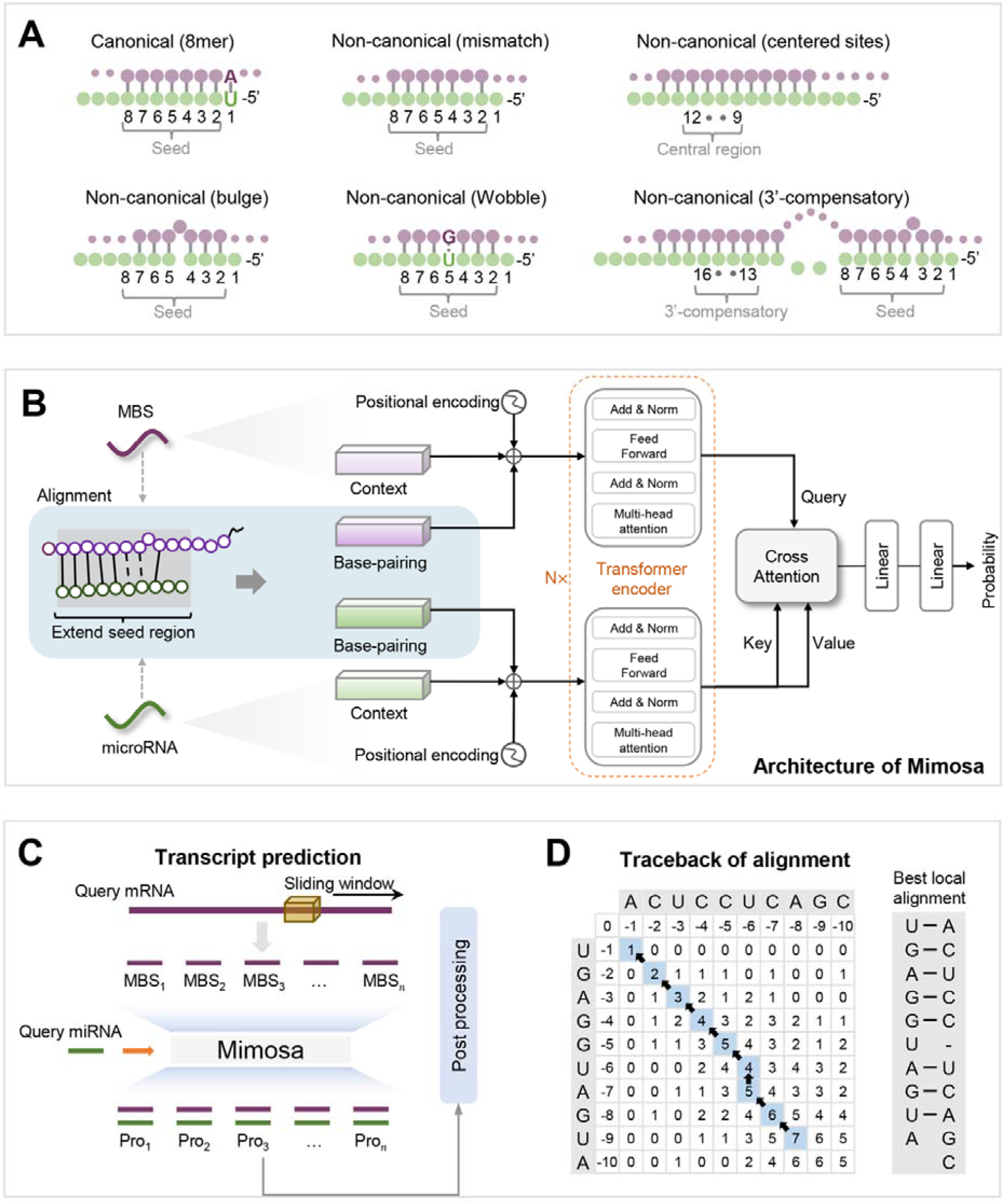
(**A**) depicts several examples of both canonical and non-canonical sites in miRNA targeting. (**B**) presents the framework of Mimosa, a dual-input model that specifically integrates base-pairing patterns into the model inputs. (**C**) outlines the gene-level prediction process for miRNA-mRNA pairs, highlighting that our model is trained on site-level samples (miRNA-MBS pairs). (**D**) illustrates the traceback process in our alignment, as well as the detailed base-pairing pattern detected.

Over the past two decades, a number of computational approaches have been developed for miRNA target identification, driven by their cost-effective and easy accessibility (19). These approaches predominantly rely on manual feature engineering for decision-making, focusing on gene-level predictions and incorporating site-level features such as seed matching (20-22). Typically, they support a limited array of seed match types, including 8mer, 7mer-m8, 7mer-A1, 6mer, and offset 6mer, all considered canonical sites. In contrast, deep learning approaches have shown a remarkable ability to automatically discern intricate data patterns compared to those reliant on feature engineering (23-26). For instance, Lee *et al*. introduced deepTarget, a model utilizing auto-encoders and stacked recurrent neural networks to analyze sequence interactions (27). Similarly, Pla *et al*. developed miRAW, a deep learning model with eight hidden layers designed for learning features and assessing functionality, with a particular focus on non-canonical sites (28). This model uses a pre-selection step for candidate target sites (CTS) to widen the range of potential miRNA targets. Subsequently, TargetNet expanded the selection criteria for CTS and leveraged the ResNet architecture to enhance the predictive performance (29). Despite the challenges in characterizing non-canonical sites due to their variable base-pairing and conservation, they are recognized for their moderate role in gene suppression, potentially due to a reduced number of AGO-miRNA complexes or weaker binding affinity (30-32). The interplay between canonical and non-canonical sites may offer a refined control over gene suppression mechanisms (33). Therefore, the emphasis on identifying non-canonical sites, as seen with miRAW and TargetNet, is crucial for advancing predictive models. However, relying solely on CTS selection could overlook biologically significant non-canonical sites that do not fit the predefined criteria, underscoring the need to strengthen models’ capabilities in predicting these sites.

In this study, we proposed Mimosa, a deep learning framework for the identification of miRNA targets that shifts away from traditional candidate target site (CTS) selection methods. The core innovation of Mimosa lies in its capacity to autonomously identify non-canonical binding sites by incorporating base-pairing patterns directly into the model’s training. This is achieved using a dynamic programming algorithm with a scoring strategy based on pairing stability, enabling the identification of optimal local alignments and the creation of base-pairing embeddings for input into the model. Such integration enables Mimosa to autonomously recognize and classify various types of non-canonical sites without manual pre-specification of pairing configurations. The foundation of Mimosa is the Transformer architecture, a robust deep learning framework renowned for its efficacy in addressing long-range dependencies in natural language processing (NLP) tasks (34). The Transformer’s ability to assimilate global information through its self-attention mechanism has not only led to remarkable successes in NLP but has also inspired a plethora of bioinformatics tools, thanks to the conceptual parallels between biological sequences and linguistic constructs (35). Mimosa harnesses this architecture to extract and integrate sequence context, positional information, and base-pairing interactions, thus functioning as a standalone and independent tool that does not rely on any third-party software. This design ensures streamlined, end-to-end predictions and optimizes the user journey. To further this goal, we have developed a user-friendly web server for Mimosa, aimed at providing easier access to advanced miRNA target identification.

**[Figure 1]**

## 2. Materials and methods

### 2.1 The benchmark dataset

This study employed a benchmark dataset compiled from experimentally validated miRNA-mRNA functional interactions (28), obtained from Diana TarBase (6) and MirTarBase (36). The dataset initially consists of 303,912 positive and 1,096 negative gene-level interaction entries for *Homo sapiens*. Site-specific binding data from PAR-Clip (37) and CLASH (38) experiments were integrated to enhance the dataset with precise interaction locations. In addition, thermodynamic properties, including minimum free energy (MFE) and species conservation, were incorporated to refine the dataset. Consequently, the benchmark is structured into a training set at the site-level, comprising 58,793 miRNA and miRNA pairing site (MBS) pairs. Each MBS consists of a 30-nt core pairing region flanked by an additional 5 nucleotides on either side. Moreover, ten test sets at the gene-level were established, each containing 548 positive and 548 negative miRNA-mRNA pairs. It is worth noting that ‘mRNA’ specifically refers to 3’UTRs, the primary region for miRNA binding. Although 5’UTRs and open reading frame (ORF) regions may facilitate some interactions, they are not the focus of the model’s development. Notably, this study adopted TargetNet’s methodology to divide the training set into a training subset, comprising 26,995 positives and 27,469 negatives, and a validation subset for parameter optimization, containing 2,193 positives and 2,136 negatives. This approach ensures consistent and comparative analysis of the model’s effectiveness.

### 2.2 The overall framework of Mimosa

To provide a clear understanding of Mimosa’s framework, we focus on a single miRNA-MBS pair, as depicted in **Figure 1B**. Mimosa is designed to process such pairs separately, requiring distinct inputs for the miRNA and MBS. The generation of these inputs initiates with the tokenization of sequences, assigning tokens for A (adenine), C (cytosine), G (guanine), U (uracil), and an additional padding token X. This standardizes the lengths of miRNA and MBS inputs to 30 and 40 tokens, respectively. Subsequently, the inputs undergo refinement through the integration of three types of embeddings: context embedding (*E*_*sequence*_), which captures contextual relevance through a high-dimensional embedding transformation; positional embedding (*E*_*position*_), initialized from a normal distribution to accurately represent the positions of nucleotides; and base-pairing embedding (*E*_*pairing*_), calculated from sequence alignment procedure to encode base pairing pattern. The inputs for miRNA and MBS are then formed by summing these embeddings, as follows:

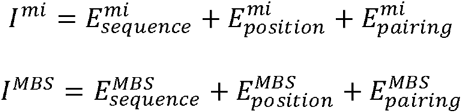

Mimosa’s model employs two separate Transformer encoder modules to process inputs and extract critical attributes from miRNA-MBS pairs. Each encoder composes multiple blocks, incorporating a self-attention mechanism and a feed-forward network for in-depth sequence analysis. Subsequently, a cross-attention layer integrates miRNA features to identify essential elements within the MBS. The results from this layer are then passed through linear layers, culminating in the final predictions for each miRNA-MBS pair. During testing, Mimosa accommodates miRNA-mRNA pairs with mRNA lengths ranging from 6-nt to 32,870-nt. As illustrated in **Figure 1C**, the model addresses this variability by employing a sliding window technique to generate subsets of longer mRNAs, consisting of 40-nt MBS segments. For mRNAs shorter than 40-nt, padding is applied. This systematic use of the sliding window and padding ensures consistent input lengths, which are crucial for the transitioning from site-level to gene-level predictions. If at least one MBS segment is predicted to form a functional pair with the miRNA, the entire mRNA is considered capable of functional interaction.

#### 2.2.1 Sequence alignment

To tackle the intricate non-canonical base-pairing patterns, our model incorporates these patterns directly into its training process. This enables the model to discern and derive rules from a diverse array of base-pairing patterns. Our approach employs the Smith-Waterman algorithm, a dynamic programming method renowned for optimal local alignment, to perform sequence alignment and extract base-pairing patterns for miRNA-MBS pairs. Widely employed in biological sequence similarity detection (39,40), this algorithm assigns similarity scores between nucleotides of two sequences. In our study, however, we have adapted the algorithm to assign scores based on the stability of base-pairing. We classify base pairings into three categories: Watson-Crick pairings (A-U and C-G), recognized for their most thermodynamic stability, are assigned a score of 1; wobble pairings (G-U), though atypical, hold functional significance and receive a score of 0; non-binding states such as gaps and mismatches are scored as -1 to indicate their instability. This alignment process involves initializing a scoring matrix and populating it based on our predefined scoring scheme (Supplementary **Algorithm 1**). A systematic traceback process then allows for the determination of the optimal pairing pattern, as depicted in **Figure 1D**. Our alignment specifically targets the extended seed region, encompassing miRNA positions 1-10 in the 5’-to-3’ direction and MBS positions 1-10 in the 3’to-5’ direction, a critical region highlighted in previous studies (28,29). This customized sequence alignment rapidly and precisely identifies base-pairing patterns, enriching the sequence characteristics and enhancing the model’s learning capability.

#### 2.2.2 The network architecture

##### (i) Transformer encoder

To effectively process sequence inputs, Mimosa integrates the Transformer encoder module into its architecture. The encoder consists of multiple identical blocks, each featuring a multi-head self-attention sub-layer as its core component, crucial for highlighting inter-token relationships. This sub-layer operates by first deriving three key matrices from the input *X*: *Q* (Query), *K* (Key), and *V* (Value), as denoted by *Q*= *XW*^*Q*^, *K*= *XW*^*K*^, and *V* = *XW*^*V*^, where *W*^*Q*^, *W*^*K*^, *W*^*V*^ are learnable parameters. The attention scores for the tokens in X are calculated using the formula 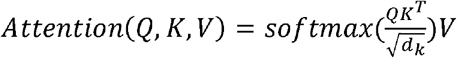, where *d*_*k*_ denotes the dimension of *K*. The term ‘multi-head’ refers to the execution of multiple such attention operations in parallel, with the fusion of these results presented in the following equation:

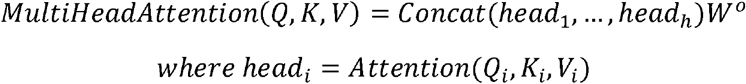

In addition to the multi-head self-attention mechanism, each block in the encoder incorporates a feed-forward network. This network consists of two linear transformations with rectified linear unit (ReLU) activation, defined as:

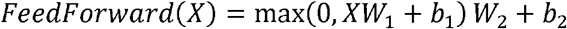

where *W*_l_, *W*_2_, *b*_l_, and *b*_2_ are learnable parameters. Following both the multi-head self-attention and the feed-forward network, a combination of residual connections and layer normalization is applied, alleviating the vanishing gradient problem and accelerating convergence. This implementation can be illustrated as follows:

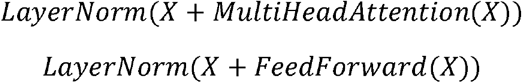

##### (ii) Cross-attention layer

The cross-attention in our model has been utilized to integrate outputs from two separate encoders. Fundamentally different from the self-attention described above, which calculates attention scores by using one data set for both the query and the key, cross-attention uniquely assigns scores by using one data set as the query and referencing a different data set as the key. This method effectively draws connections between different data sources, thereby enhancing the overall performance of the model. The specifics of this operation are defined as follows:

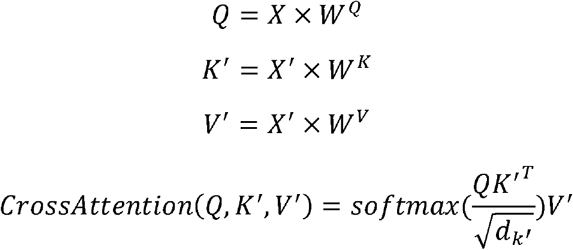

Here, *X* denotes the query input and *X*, represents reference input, with the cross-attention output dimensionally consistent with *X*.

#### 2.2.3 Training and evaluation strategies

In this study, we manually fine-tuned Mimosa’s hyperparameters to enhance its performance. The embedding dimensions for *E*_*sequence*_, *E*_*position*_, and *E*_*pairing*_ were set to 64. The encoder was structured with 16 blocks, each featuring 8-head self-attention. A similar design was adopted for the cross-attention layer, which consists of 16 blocks with 8 heads each. The training parameters include a learning rate of 1e-4, a batch size of 256, and the Adam optimization algorithm with 1e-5 weight decay. To avoid overfitting, a dropout rate of 0.1 was integrated into each encoder module. For loss calculation, we applied binary cross-entropy, represented by the formula:

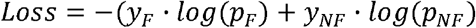

where *y*_*F*_ and *y*_*NF*_ represent the truth labels for functional and non-functional, respectively, and *p*_*F*_ and *p*_*NF*_ are their prediction probabilities. The model underwent 40 epochs of training, with the version exhibiting the minimum loss selected as the final model.

We evaluated the model using six commonly employed metrics, derived from the counts of true positives (TP), true negatives (TN), false positives (FN), and false positives (FP). These metrics include accuracy, F1 score, recall, specificity, positive predictive value (PPV), and negative predictive value (NPV). The definitions of these metrics are as follows: accuracy is calculated as (*TP* + *TN*)/(*TP* + *FP* + *TN* + *FN*), F1 score is calculated as 2*TP*/(2*TP* + *FP* + *FN*), recall as *TP*/(*TP* + *FN*), specificity as *TN*/(*TN*+ *FP*), PPV as *TP*/(*TP* + *FP*), and NPV as *TN*/(*FN* + *TN*).

## 3. Results and discussion

### 3.1 In-depth analysis of model architecture

This section delves into the architecture of Mimosa, highlighting its pivotal components and strategies across four domains: the integration of base-pairing patterns, the formulation of a base-pairing detection strategy, the selection of input processing feature extractors, and the amalgamation of these extracted features. Initially, our approach of solely utilizing conventional contextual and positional encoding for input data did not facilitate model convergence, a challenge we attributed to the compact data size and the brevity of sequences involved. The incorporation of base-pairing patterns notably enhanced model performance, leading to a 16.22% increase in accuracy and a 9.65% improvement in F1 score (refer to **Figure 2A**). We explored three distinct scoring strategies for detecting base-pairings: (1) assigning a score of 1 to both Watson-Crick and wobble pairings, while treating gaps and mismatches with a score of -1; (2) scoring Watson-Crick and wobble pairing as 1, with gaps and mismatches scored as 0; (3) attributing a score of 1 to Watson-Crick pairings, 0 to wobble pairings, and -1 to gaps and mismatches. The scoring system that most accurately reflected interaction stability delivered substantial performance enhancements, improving accuracy by a minimum of 5.89% and the F1 score by 1.8% (shown in **Figure 2B**). In our comparison of input processing mechanisms, we scrutinized the complete Transformer model (Transformer-complete), its encoder component (Transformer-encoder), and Bert, an advanced NLP model predicated on the Transformer encoder (41). The Transformer-encoder emerged as the optimal choice due to its superior accuracy, F1 score, and NPV, as depicted in **Figure 2C**. During the feature integration phase, employing cross-attention markedly outperformed simple tensor concatenation, specifically demonstrating a 7% accuracy increment. This finding positioned cross-attention as a cornerstone of Mimosa’s architectural framework (illustrated in **Figure 2D**).

**Figure 2.**
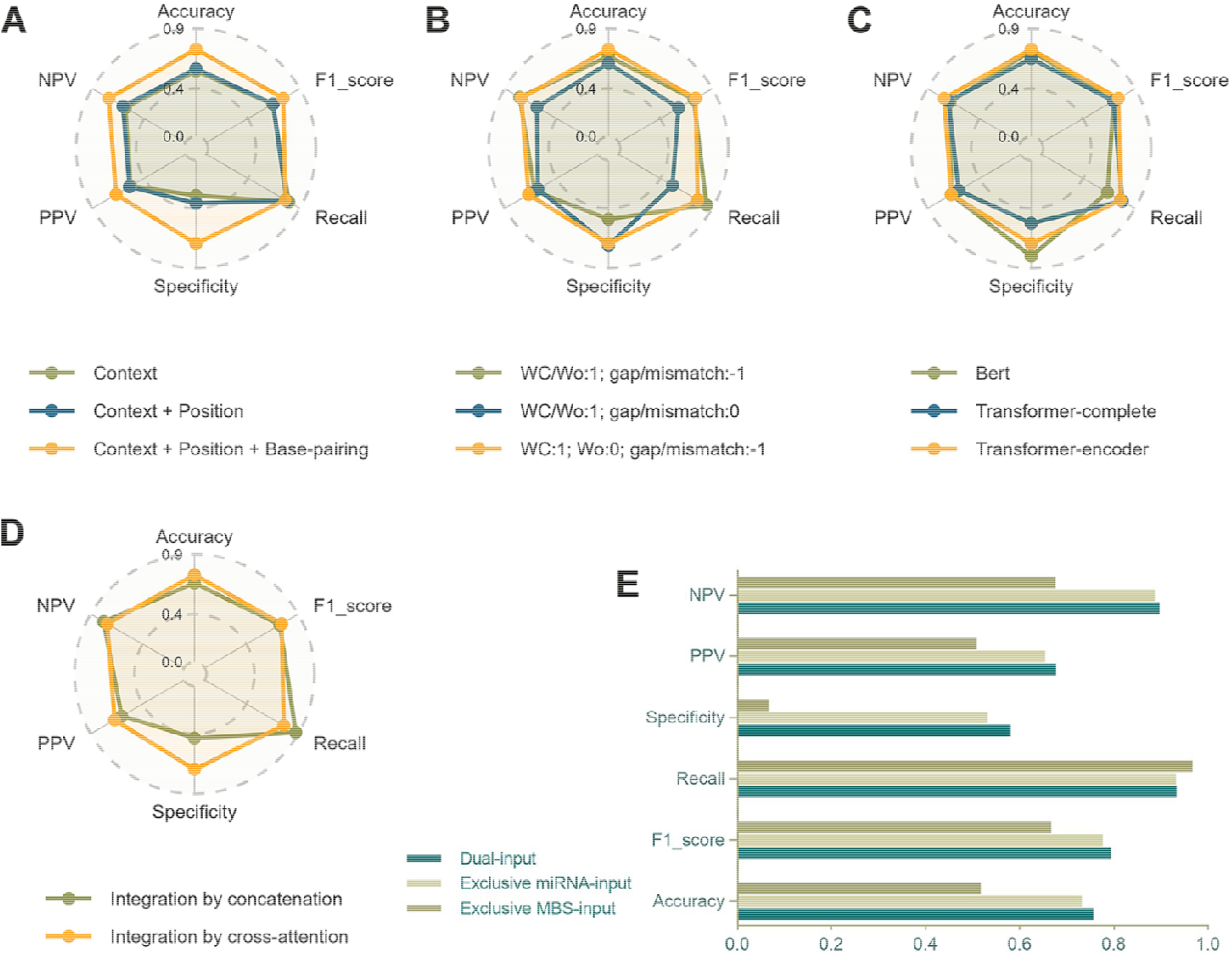
Analysis of Mimosa’s model architecture components. **(A)** Illustrates the impact of different input encodings, underscoring the critical role of integrating base-pairing patterns for enhanced model performance. **(B)** Analyzes three distinct scoring strategies for base-pairing patterns, categorizing them into Watson-Crick (WC) pairings and Wobble (Wo) pairing, to determine the most effective approach. **(C)** Evaluates three different structures for processing inputs, with the Transformer-encoder emerging as the superior choice for accuracy and performance. **(D)** Explores various techniques for merging outputs from the miRNA and MBS Transformer-encoders, demonstrating the effectiveness of cross-attention mechanisms. **(E)** Describes an ablation study aimed at evaluating the comparative advantages of dual-input versus single-input models in terms of predictive accuracy and generalization.

To ascertain whether the integration of dual inputs is imperative for our model, and to evaluate the feasibility of transitioning to a single-input system, we conducted an ablation study. This study entailed retraining our model separately with only miRNA inputs and then only MBS inputs, to compare the individual performance against the dual-input setup. Results from this investigation indicated that the model trained solely with MBS inputs yielded superior performance relative to the dual-input structure during the training phase, as evidenced by the results in **Supplementary Table S1**. However, this superior performance was not sustained into the testing phase, a discrepancy highlighted in **Figure 2E**. This discrepancy is attributed to potential overfitting during the training phase, which seemed to compromise the model’s generalization ability to unseen data. Conversely, the dual-input model displayed steady and uniform predictive performance across both the validation and test datasets. This consistency underscores the model’s robust generalization capability and affirms the architectural superiority of the dual-input model.

**[Figure 2]**

### 3.2 Performance evaluation of Mimosa

Given that 99.38% of mRNA sequences in our test sets exceeded 40 nucleotides, the selection of an appropriate step size for the sliding window was critical. The determination of Mimosa’s default step size was based on a thorough analysis of its impact on performance. Our experiments involved exploring step sizes ranging from 1 to 40 across all test sets. As illustrated in **Figure 3A**, accuracy fluctuated between 0.7354 and 0.7865, and the F1 score ranged from 0.7633 to 0.8109, indicating a marginal difference of approximately 5% for both metrics. We observed a slight positive trend in both accuracy and F1 score with step sizes smaller than 6, transitioning to a more irregular pattern at larger step sizes. Although the precise reasons for this variability remain unclear, it is evident that increasing step size results in less stable model performance. To facilitate comparison and optimize computation efficiency, all comparative experiments in this study utilized a step size of 5, unless otherwise specified.

**Figure 3.**
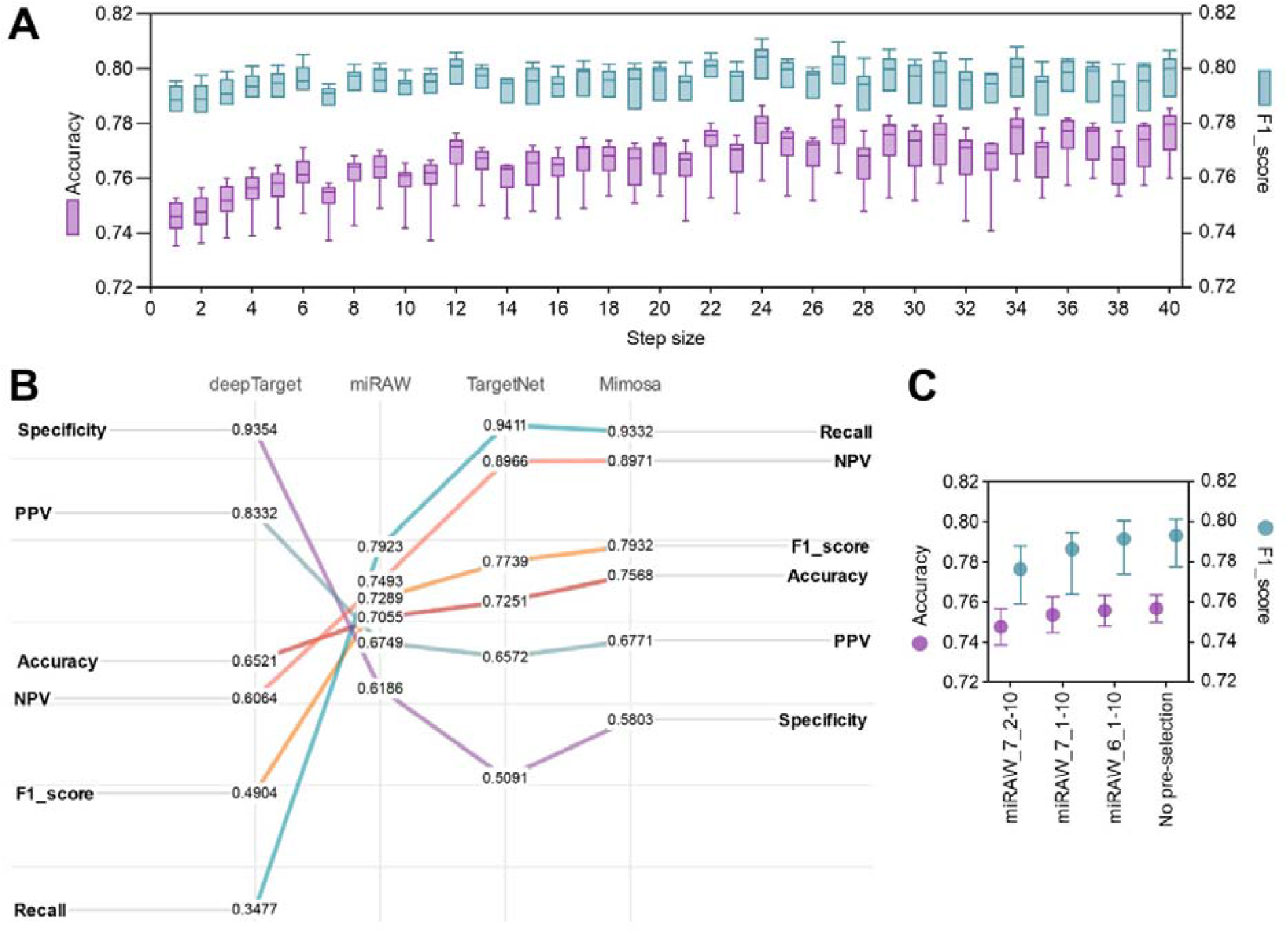
Evaluation of Mimosa’s predictive performance. **(A)** Analysis of step-size variation: this min-to-max boxplot illustrates the fluctuations in accuracy and F1 score across ten test sets, highlighting the impact of step size on the sliding window technique. **(B)** Model comparison: comparison of Mimosa’s metrics with other deep learning models. **(C)** Candidate target site (CTS) selection analysis: evaluation of the effect of different CTS criteria on Mimosa’s accuracy and F1 score.

To assess Mimosa’s effectiveness, we conducted a comparative analysis against state-of-the-art deep learning approaches, including deepTarget, miRAW, and TargetNet. Utilizing data for a previous study (29), we examined the average performance across ten test sets for each model. As depicted in **Figure 3B**, deepTarget displayed significant variability in its metrics, exhibiting the highest specificity but the lowest recall. Conversely, miRAW demonstrated the least variation in metrics, albeit with lower accuracy and F1 score compared to TargetNet. Mimosa surpassed TargetNet in most metrics, although with a slightly weaker recall. Notably, Mimosa achieved the highest NPV, F1 score and accuracy among the four models, highlighting its superior predictive capabilities. To ensure equitable comparison with TargetNet, Mimosa’s performance was also assessed using a step size of 1, consistent with TargetNet’s methodology (refere to **Supplementary Table S2**). Despite a minor decrease in F1 score and accuracy, Mimosa consistently outperformed TargetNet across all assessed metrics.

It is reasonable to deduce that Mimosa’s intricate base-pairing analysis obviates the necessity for pre-selecting candidate target sites (CTS) prior to prediction. To verify this assertion, we investigated the impact of employing CTS selection on performance. Specifically, we applied three CTS selection criteria outlined in [24] and compared them against the scenario without CTS selection. These criteria included: (1) miRAW-6-1:10, necessitating a minimum of 6 base pairs within positions 1-10; (2) miRAW-7-1:10, requiring at least 7 base pairs within positions 1-10; and (3) miRAW-7-2:10, mandating a minimum of 7 base pairs within positions 2-10, encompassing both Watson-Crick and wobble pairings. Comparative results presented in **Figure 3C** illustrate that while the selection criteria marginally impact accuracy and F1 score, stricter criteria notably diminish these metrics, especially when compared to the scenario without pre-selection criteria. This decline may stem from inadvertently filtering out viable miRNA targets during the pre-selection process. In summary, Mimosa’s efficacy remains unaffected by CTS selection criteria.

**[Figure 3]**

### 3.3 Effect of binding free energy

The binding free energy (Δ*G*) between miRNA and its target sites is a crucial thermodynamic parameter used to assess the stability of double-stranded molecular structures, where lower Δ *G* values indicate more stable bindings. Δ *G* has been widely employed as a key filtering criterion in various established approaches, typically employing a threshold between -17 and -12 kcal/mol (19). To explore the influence of Δ*G* filtering on the Mimosa model, we embarked on a thorough re-assessment. This re-assessment entailed the extension of our functional determination criteria to include not only a predictive probability exceeding 0.5 but also a Δ*G* value below a specified threshold. This threshold underwent rigorous testing across a range from -20 to -1 kcal/mol. Δ *G* calculations were performed using the RNAduplex software (42). **Figure 4A** illustrates how accuracy and F1 score vary with changes in Δ *G*, noting that a Δ*G* value of 0 corresponds to Mimosa’s baseline prediction without Δ*G* filtering. We observed a significant decline in both accuracy and F1 score as the Δ*G* value became less negative, suggesting that a more stringent criterion reduces the identification of functional targets. Additionally, we found that Δ *G* thresholds between -10 to 0 had minimal impact on our test results, possibly due to our evaluation strategy relying on identifying at least one site-level target to define the function of the entire sequence. To further investigate changes in site-level targets, we randomly selected three miRNA-mRNA pairs and analyzed the variation in the number of site-level targets with Δ*G*, as shown in **Figure 4B**. It became evident that Δ*G* filtering reduced the number of predicted site-level targets, potentially decreasing the high false-positive rates associated with sequence-based predictions. Although our Mimosa model excludes Δ*G* filtering, combining predictions from multiple perspectives still holds promise.

**Figure 4.**
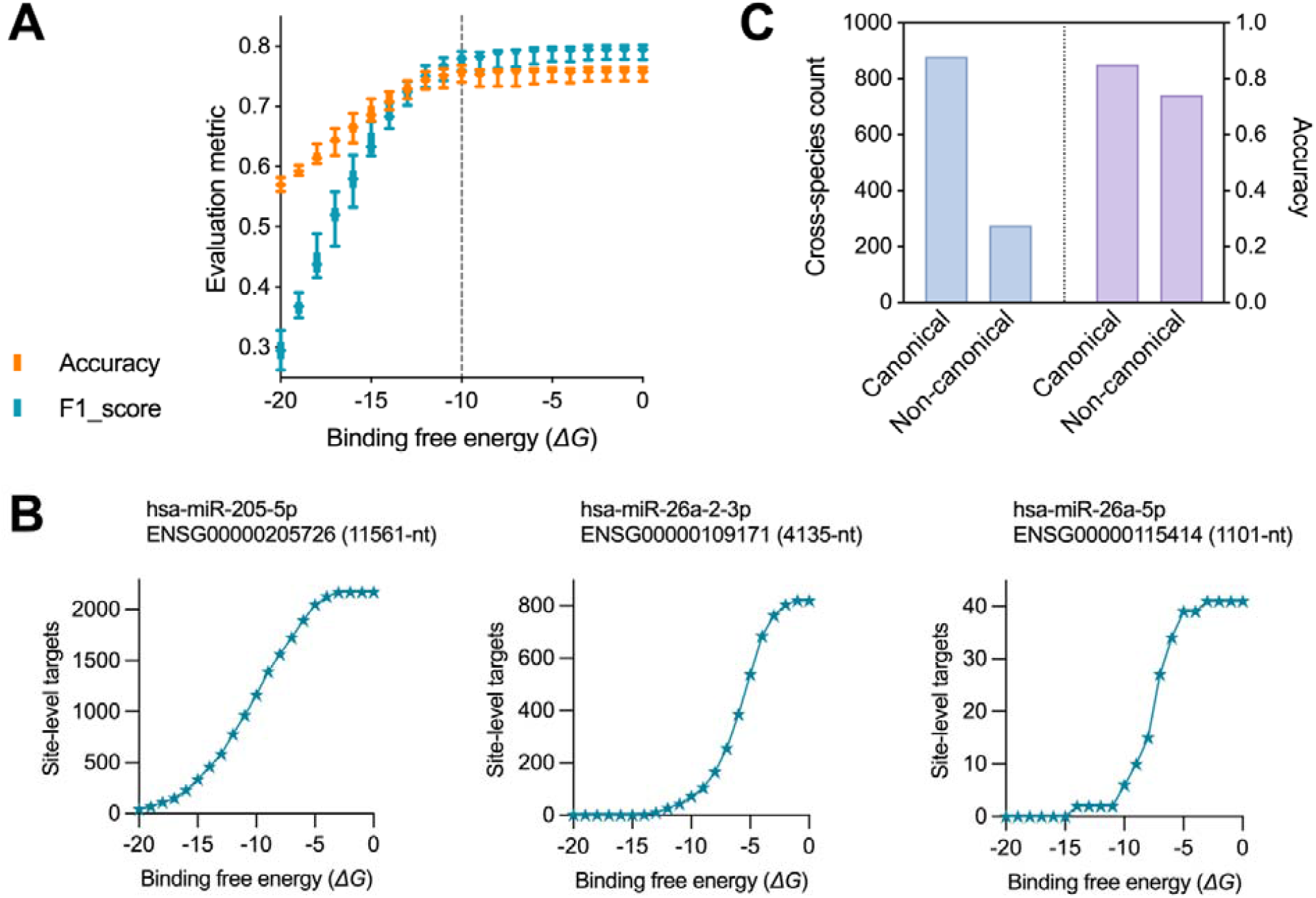
**(A)** illustrates the impact of binding free energy (Δ*G*) on accuracy and F1 score (average results of ten test sets), highlighting Δ*G* ‘s influence on gene-level predictions. **(B)** reveals the variation in the count of predicted target sites in three cases, showing Δ*G*’s influence on site-level predictions. **(C)** showcases Mimosa’s overall accuracy for both canonical and non-canonical sites in predictions for non-human species.

**[Figure 4]**

### 3.4 Mimosa’s prediction across different species

Considering the conservation of miRNA regulatory mechanisms across species, we evaluated the efficacy of Mimosa in predicting miRNA targets in non-human organisms. We curated a dataset comprising 1156 experimentally validated miRNA-target pairs from 12 cellular organisms sourced from the MirTarBase database (43). Our analysis revealed that Mimosa demonstrated commendable performance by accurately predicting 955 out of the 1156 pairs, resulting in an accuracy rate of 82.61%. We further categorized these targets into canonical and non-canonical categories for a more nuanced analysis. Canonical sites encompassed patterns such as 8mer, 8mer-A1, 7mer-m8, 7mer-A1, 6mer (p1-p6), 6mer, and 6mer (p3-p8) (detailed in **Table S3**), with pairs not conforming to these patterns classified as non-canonical. The statistical outcomes of Mimosa’s predictions for each species are presented in **Table 1**. Of the total 880 canonical pairs, Mimosa successfully predicted 750, achieving an accuracy of 85.23%. Despite the relative scarcity of non-canonical pairs, totaling 276, Mimosa maintained a high accuracy of 74.28%, identifying 205 of these pairs, as illustrated in **Figure 4C**. We infer that the slightly lower performance at non-canonical sites may be attributed to the limited representation of non-canonical patterns in the training dataset, thereby constraining the model’s ability to learn a broader range of patterns. Nevertheless, these overall results highlight Mimosa’s broad applicability and reliability, positioning it as a valuable tool for predicting miRNA targets across a diverse species.

**[Table 1]**

### 3.5 Operating Mimosa

To enhance accessibility for the scientific community, we have made available a user-friendly webserver for Mimosa (http://monash.bioweb.cloud.edu.au/Mimosa/) dedicated to binary interaction predictions for miRNA-mRNA pairs. This webserver accommodates the processing of up to ten pairs simultaneously, allowing sequences of up to 8,000-nt in length. Its interface provides two separate textboxes for submitting miRNA and mRNA sequences in FASTA format, ensuring each pair is matched correctly. Furthermore, our platform offers specialized functionality targeting the 3’UTR region within complete mRNA sequences. Users can designate ‘3’UTR’ as the predictive region from a dropdown menu, enabling the system to identify and utilize the longest 3’UTR segment for prediction. To optimize usability, we have set the default step size of the sliding window at five, though users have the option to customize this setting according to their preferences. The results page delivers a binary predictive label for each query pair, indicating whether the interaction is functional or non-functional. For users interested in delving in-depth information on target sites and specific base-pairing patterns, a link to our code repository is provided for further exploration at: https://github.com/biyueeee/Mimosa.

### 3.6 Exploring miR-BART6-5p Targeting Dicer: A Case Study

Epstein-Barr virus (EBV) is a gamma-herpesvirus associated with various lymphoid and epithelial malignancies (44). It infects over 90% of the global adult population, persisting as a lifelong infection. Studies indicate that EBV-infected cells express abundant EBV miRNAs, with more than 40 mature EBV miRNAs identified to date (45). Research by Iizasa *et al*. highlighted the role of one such miRNA, ebv-miR-BART6-5p (5’-UAAGGUUGGUCCAAUCCAUAGG-3’), in targeting the 3’UTR of human Dicer mRNA, thereby suppressing the expression of miRNAs from both viral and host origins (46). This suppression is critical for viral persistence and latent infection sustainment. In this section, we utilized Mimosa to analyse the targeting of ebv-miR-BART6-5p on the Dicer mRNA 3’UTR. Mimosa identified 191 functional segments within the Dicer 3’UTR using a size step of 1. These segments were evenly distributed across the Dicer 3’UTR, with predominantly non-canonical and only twelve canonical segments. As depicted in **Figure 5**, our analysis revealed a shared non-canonical site within a target-dense interval, featuring a wobble pairing and a bulge, suggesting potential targeting by ebv-miR-BART6-5p. Among the twelve canonical functional segments, we identified seven non-overlapping target sites (Please refer to **Supplementary Table S4** for more details). Remarkably, four of the seven canonical sites identified by Mimosa have been validated as unique to the human Dicer 3’UTR using the Luciferase Reporter technology (46), exhibiting strong interaction characteristics, particularly with 7mer-A1 types. The remaining three sites display weaker interactions, characterized by 6mer and offset 6mer types. Overall, Mimosa demonstrates the comprehensive predictive capability for both canonical and non-canonical target sites of ebv-miR-BART6-5p on the Dicer 3’UTR. This analysis provides valuable insights to support and enhance research into EBV miRNA functions.

**Figure 5.**
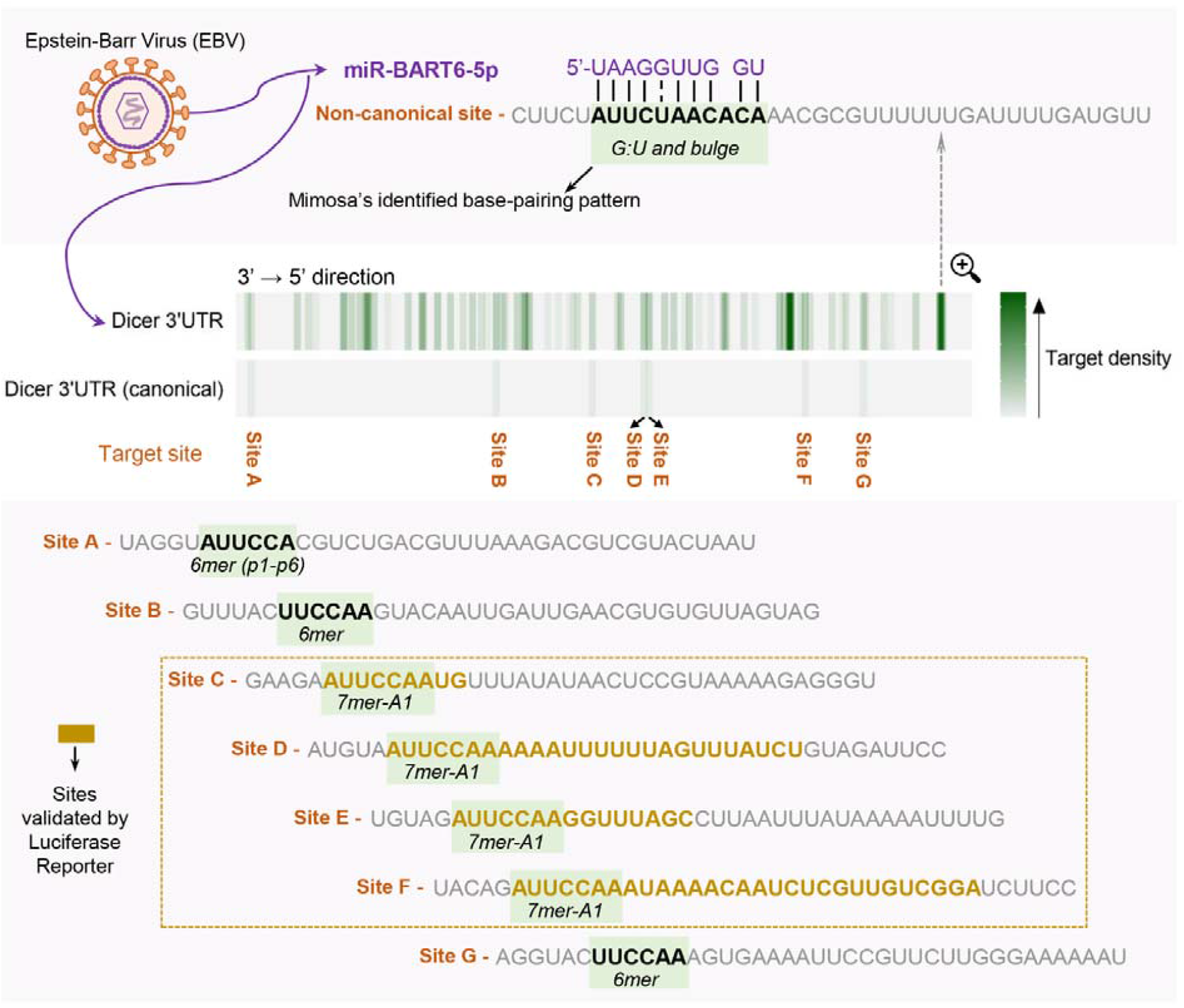
Predicted ebv-miR-BART6-5p (a miRNA of the Epstein-Barr virus) target sites by Mimosa within the Dicer mRNA 3’UTR, meticulously highlights a non-canonical site alongside all seven canonical sites.

**[Figure 5]**

## Conclusion

MicroRNAs (miRNAs) serve as pivotal regulatory factors in numerous cellular processes, exerting significant influence on cell differentiation, development, and homeostasis. Identifying miRNA targets is fundamental for unravelling the functions and mechanisms of these regulatory molecules. To expedite this process, we introduced Mimosa, a Transformer-based method designed for predicting miRNA targets. A notable strength of Mimosa lies in its consideration of base-pairing patterns, which not only broadens its recognition scope to encompass both canonical and non-canonical sites but also reduces reliance on pre-selecting candidate targets during prediction. By doing so, Mimosa enables the detection of non-canonical sites that might otherwise be overlooked due to stringent pre-selection criteria. Our evaluations demonstrate that Mimosa outperforms existing deep learning models and effectively identifies miRNA targets across diverse species. Meanwhile, the user-friendly interface of Mimosa’s webserver provides a streamlined and efficient means of pinpointing target sites accurately, thereby enhancing the analysis process. We anticipate that Mimosa will emerge as a vital tool for researchers investigating miRNA-specific targets and exploring the intricate small RNA regulatory network.

It is important to acknowledge that the predictions generated by Mimosa for non-canonical sites may exhibit a higher rate of ‘false positives’, as evidenced in our case study. MiRNA regulation is governed by a complex interplay of multiple factors, including miRNA expression levels, the assembly and activity of the miRNA-induced silencing complex (miRISC), and the modulation of competing endogenous RNAs (ceRNAs). This intricate network enables miRNAs to finely tune gene expression. The target sites identified by Mimosa reflect potential regulatory mechanisms based on genetic predisposition, with non-canonical sites poised within genes, awaiting activation under appropriate conditions, such as heightened miRNA expression. However, the activation of these sites depends not only on miRNA presence but also on the synergy of other factors, including specific cellular states and external signals. Furthermore, the inherent instability of non-canonical sites complicates their detection and validation. Therefore, while Mimosa offers a predictive framework, comprehensively considering all relevant factors and their dynamic interplay remains a challenge in understanding the full complexity of miRNA interactions.

### Key points

- Comprehensive identification of miRNA targets is crucial for deepening our understanding of the mechanisms underlying the precise regulation of gene expression.
- We introduce Mimosa, a deep learning-based approach for miRNA target identification, adept at recognizing the elusive ‘non-canonical sites’ in human species.
- Mimosa demonstrates superior performance compared to existing deep learning models and also exhibits excellent generalization capacity across a variety of non-human species.
- Case studies of Mimosa of miR-BART6-5p targeting Dicer provide a useful reference for target sites, offering insight into the research related to EBV miRNA functions.

