## Supplementary Material for "Advancing microRNA Target Site Prediction with Transformer and Base-Pairing Patterns"

Supplementary Pseudocode and Tables

**Algorithm S1**. The pseudocode for the sequence alignment algorithm applied in the Mimosa model.

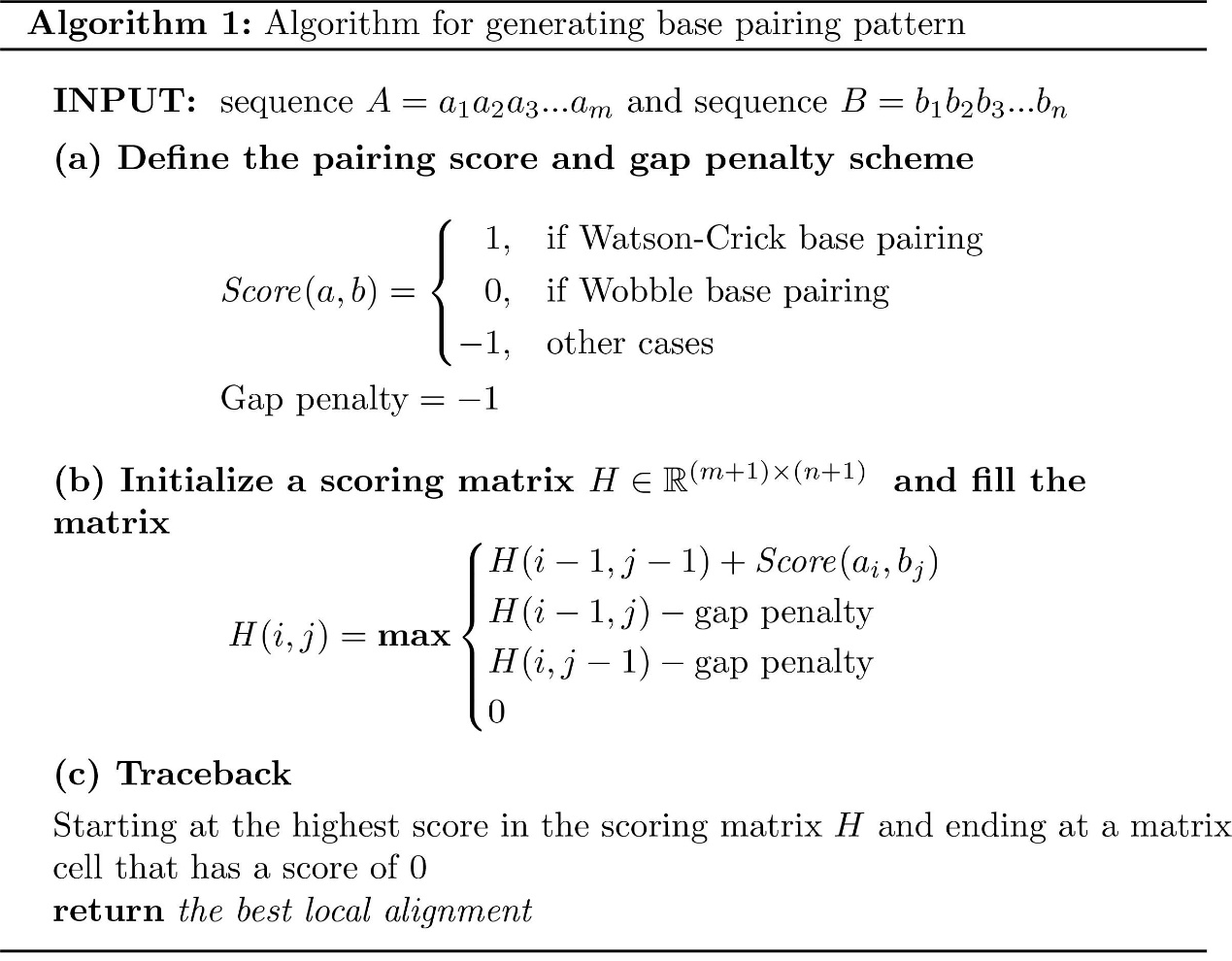

**Table S1**. Ablation study on the impact of dual-input vs. single-input configurations on model performance.

| Model | F1 score | Accuracy | Recall | Specificity | PPV | NPV |
| --- | --- | --- | --- | --- | --- | --- |
| Dual-input | 0.7400 | 0.7293 | 0.7606 | 0.6971 | 0.6693 | 0.7393 |
| Exclusive miRNA-input | 0.6876 | 0.6639 | 0.7301 | 0.5960 | 0.6111 | 0.6826 |
| Exclusive MBS-input | 0.7634 | 0.7741 | 0.7196 | 0.8301 | 0.7271 | 0.7425 |

**Note**: This comparison focuses on model training, specifically the results obtained from the validation set.

**Table S2**. Comparative analysis of deep learning models across ten test sets' average results.

| Method | F1 score | Accuracy | Recall | Specificity | PPV | NPV |
| --- | --- | --- | --- | --- | --- | --- |
| deepTarget | 0.4904 | 0.6521 | 0.3477 | 0.9354 | 0.8332 | 0.6064 |
| miRAW | 0.7289 | 0.7055 | 0.7923 | 0.6186 | 0.6749 | 0.7493 |
| TargetNet | 0.7739 | 0.7251 | 0.9411 | 0.5091 | 0.6572 | 0.8966 |
| Mimosa (CTS: miRAW-7-2:10) | 0.7765 | 0.7477 | 0.8768 | 0.6186 | 0.6727 | 0.8344 |
| Mimosa (CTS: miRAW-7-1:10) | 0.7864 | 0.7537 | 0.9071 | 0.6004 | 0.6762 | 0.8665 |
| Mimosa (CTS: miRAW-6-1:10) | 0.7915 | 0.7557 | 0.9276 | 0.5839 | 0.6766 | 0.8901 |
| Mimosa (Size step=5) | 0.7932 | 0.7568 | 0.9332 | 0.5803 | 0.6771 | 0.8971 |
| Mimosa (Size step=1) | 0.7884 | 0.7457 | 0.9476 | 0.5438 | 0.6659 | 0.9124 |

**Table S3**. The types of canonical sites and their definitions.

| Seed match type | Definition |
| --- | --- |
| 8mer | p1-p8 Watson-Crick match |
| 8mer-A1 | p2-p8 Watson-Crick match with ‘A’ at p1 |
| 7mer-m8 | p2-p8 Watson-Crick match |
| 7mer-A1 | p2-p7 Watson-Crick match with ‘A’ at p1 |
| 6mer (p1-p6) | p1-p6 Watson-Crick match |
| 6mer | p2-p7 Watson-Crick match |
| 6mer (p3-p8) | p3-p8 Watson-Crick match |

**Table S4**. Comprehensive list of miR-BART6-5p functional target sites within the Dicer 3’UTR identified by Mimosa.

| Index | Target site (3’→5’ direction) | Base-pairing pattern |
| --- | --- | --- |
| 1 | UGUUGAUUUCCAUUAACACUAGGUAUUCCACGUCUGACGU | UAA-GGU  \|\|\|-\|\|\|  AUUUCCA |
| 2 | GUUGAUUUCCAUUAACACUAGGUAUUCCACGUCUGACGUU | AAGGU  \|\|\|\|\|  UUCCA |
| 3 | AAAAUUUUUCCAUUUAAUUCUCCAUAAUAUCAAUGUCACG |  |
| 4 | AAAUUUUUCCAUUUAAUUCUCCAUAAUAUCAAUGUCACGU |  |
| 5 | UAGGUAUUCCACGUCUGACGUUUAAAGACGUCGUACUAAU | UAAGGU  \|\|\|\|\|\|  AUUCCA |
| 6 | AGGUAUUCCACGUCUGACGUUUAAAGACGUCGUACUAAUG | AGGU  \|\|\|\|  UCCA |
| 7 | AAUUUUUCCAUUUAAUUCUCCAUAAUAUCAAUGUCACGUU |  |
| 8 | UUUAAUUCUCCAUAAUAUCAAUGUCACGUUUUUUUUUUUU |  |
| 9 | UUAAUUCUCCAUAAUAUCAAUGUCACGUUUUUUUUUUUUU |  |
| 10 | UAAUUCUCCAUAAUAUCAAUGUCACGUUUUUUUUUUUUUA |  |
| 11 | UCUUGACUCCAUAAAUUCACUUCUCACGGAUGUUCAGAUU |  |
| 12 | CUUGACUCCAUAAAUUCACUUCUCACGGAUGUUCAGAUUA |  |
| 13 | GAAACCAUCCACUUUAUACGUACGUUGUAAUACUUUUCAU |  |
| 14 | UGGUAUAUCCAUUAUUUGUACUGUACUCAAAGAGAAGACG |  |
| 15 | CACAAUCUCCAGUAAGAUUUAAAGUAGUGAAGUGCAUUAU |  |
| 16 | ACAAUCUCCAGUAAGAUUUAAAGUAGUGAAGUGCAUUAUA |  |
| 17 | AUGAUAGUCCACGUUGUCGCUGGCCCUUGUAGUGGAAUGU |  |
| 18 | UGAUAGUCCACGUUGUCGCUGGCCCUUGUAGUGGAAUGUG |  |
| 19 | AACAGACACCAACUACAGGUUUAUUUUUACCUGAUCUUUU | GGUUGGU  \|\|\|\|\|:\|  CCAACUA |
| 20 | ACAGACACCAACUACAGGUUUAUUUUUACCUGAUCUUUUU |  |
| 21 | CAGACACCAACUACAGGUUUAUUUUUACCUGAUCUUUUUU |  |
| 22 | UAAAUGUUCAAUCAACACUUUUUAGUAAAAGUGACGAUCA | AAGGUUGGU  \|\|\|-\|\|:\|\|  UUC-AAUCA |
| 23 | UUCAAUCAACACUUUUUAGUAAAAGUGACGAUCACCUUUU | GUUG  \|\|\|\|  CAAC |
| 24 | UAAAUUCAACACGCUCAUAUACUGUGUAACGUCCGUAAUA |  |
| 25 | AGACGUCAACGAAAAAGUUCUGUGCGGAGGGGUCAGGAAA |  |
| 26 | AUCACUCAACAUUCAUAGUGCGACAGAGUUGCAGAUUACG |  |
| 27 | UUUAUGUUACCAGAGUUACGUUUAAAGGAUCUAUAACGAU | AA-GGU  \|\|-\|\|\|  UUACCA |
| 28 | GACUUUUCAUUAGUCACCAACCUUUUUAUAUUAAUAAGUU | GGUUGG  \|\|\|\|\|\|  CCAACC |
| 29 | CAUUAGUCACCAACCUUUUUAUAUUAAUAAGUUUACAUUA |  |
| 30 | AUUAGUCACCAACCUUUUUAUAUUAAUAAGUUUACAUUAA |  |
| 31 | UUAGUCACCAACCUUUUUAUAUUAAUAAGUUUACAUUAAU |  |
| 32 | UAGUCACCAACCUUUUUAUAUUAAUAAGUUUACAUUAAUU |  |
| 33 | AGUCACCAACCUUUUUAUAUUAAUAAGUUUACAUUAAUUU | GUUGG  \|\|\|\|\|  CAACC |
| 34 | UUGGUAUAUCCAUUAUUUGUACUGUACUCAAAGAGAAGAC | UA-AGGU  \|\|-\|\|\|\|  AUAUCCA |
| 35 | GGUAUAUCCAUUAUUUGUACUGUACUCAAAGAGAAGACGU | UAAGGU  \|\|-\|\|\|  AU-CCA |
| 36 | AAACCAUCCACUUUAUACGUACGUUGUAAUACUUUUCAUA |  |
| 37 | UUACGUUUACCCAAUUUCUGAGAAUUGUAUUAAAGUCUAC | AA-GG-UU  \|\|-\|\|-\|\|  UUACCCAA |
| 38 | CGUUUACCCAAUUUCUGAGAAUUGUAUUAAAGUCUACGUC | GGUU  \|\|\|\|  CCAA |
| 39 | GUUUACCCAAUUUCUGAGAAUUGUAUUAAAGUCUACGUCA |  |
| 40 | UUACUUCCAAGUACAAUUGAUUGAACGUGUGUUAGUAGUA |  |
| 41 | GACCAUCCAAAAGGUAUAAUGUAAGCCAGAGUGUAACGUU |  |
| 42 | AGAAUUCCAAUGUUUAUAUAACUCCGUAAAAAGAGGGUGA |  |
| 43 | GUAAUUCCAAAAAAUUUUUUAGUUUAUCUGUAGAUUCCAA |  |
| 44 | UAGAUUCCAAGGUUUAGCCUUAAUUUAUAAAAAUUUUGUU |  |
| 45 | GUGAGUCCCAAGGAAUCUUUCGUCUCGACUGUCAAAAGAC |  |
| 46 | UGAGUCCCAAGGAAUCUUUCGUCUCGACUGUCAAAAGACA |  |
| 47 | CAGAUUCCAAAUAAAACAAUCUCGUUGUCGGAUCUUCCGU |  |
| 48 | UAUAAGACCAAAAAUUUUACAGAAGUCAUAUGUAUACUGU |  |
| 49 | AUAAGACCAAAAAUUUUACAGAAGUCAUAUGUAUACUGUC |  |
| 50 | CGACCUCCAAAUUCGAGGUACCGGAAGAUUGUGAUUCGUU |  |
| 51 | CUACGUCAAUUUUUUUCCAUAGUUCCAGAGUCAAACCACC | AAGGU-U-GGU  \|\|\|\|\|-\|-:\|\|  UUCCAGAGUCA |
| 52 | UACGUCAAUUUUUUUCCAUAGUUCCAGAGUCAAACCACCG |  |
| 53 | UCAAUUUUUUUCCAUAGUUCCAGAGUCAAACCACCGAAGU |  |
| 54 | AAUUUUUUUCCAUAGUUCCAGAGUCAAACCACCGAAGUUA |  |
| 55 | AUUUUUUUCCAUAGUUCCAGAGUCAAACCACCGAAGUUAG |  |
| 56 | UUUUUUUCCAUAGUUCCAGAGUCAAACCACCGAAGUUAGA |  |
| 57 | UUUUUUCCAUAGUUCCAGAGUCAAACCACCGAAGUUAGAA |  |
| 58 | UUCCAUAGUUCCAGAGUCAAACCACCGAAGUUAGAACACA |  |
| 59 | UCCAUAGUUCCAGAGUCAAACCACCGAAGUUAGAACACAU |  |
| 60 | CCAUAGUUCCAGAGUCAAACCACCGAAGUUAGAACACAUU |  |
| 61 | GAACACAUUUCCCUAAUCUGUGGGAUUGUCUCGUUCUAGG | AAGG--UUGG  \|\|\|\|--\|\|:\|  UUCCCUAAUC |
| 62 | ACACAUUUCCCUAAUCUGUGGGAUUGUCUCGUUCUAGGUU |  |
| 63 | AACACAUUUCCCUAAUCUGUGGGAUUGUCUCGUUCUAGGU | UAA-GG--UUGG  \|\|\|-\|\|--\|\|:\|  AUUUCCCUAAUC |
| 64 | GCUAGACCUAAGGUCACUAGGAGACGUCACGGGUGGACGG | GG-UU  \|\|-\|\|  CCUAA |
| 65 | ACAACAGUUCCGUCCCAUAGUCUUAGAAAGACUCCCGUUU | AAGGUUGG  \|\|\|\|:-\|\|  UUCCGUCC |
| 66 | CAACAGUUCCGUCCCAUAGUCUUAGAAAGACUCCCGUUUC |  |
| 67 | AGUUCCGUCCCAUAGUCUUAGAAAGACUCCCGUUUCAUAA | AGG  \|\|\|  UCC |
| 68 | CAUUGGUCCCCUAGUGUCUUACGAAGUUUGACACUCAGUU |  |
| 69 | UUAAUGUCCUCACAAUCUCCAGUAAGAUUUAAAGUAGUGA | AG-GUUGG  \|\|-\|\|\|:\|  UCACAAUC |
| 70 | UAAUGUCCUCACAAUCUCCAGUAAGAUUUAAAGUAGUGAA |  |
| 71 | AAUGUCCUCACAAUCUCCAGUAAGAUUUAAAGUAGUGAAG |  |
| 72 | AUGUCCUCACAAUCUCCAGUAAGAUUUAAAGUAGUGAAGU |  |
| 73 | UCACAAUCUCCAGUAAGAUUUAAAGUAGUGAAGUGCAUUA | UA-AGGU  \|\|-\|\|\|\|  AUCUCCA |
| 74 | UGUAUUAAUUCUCAAGUCAAUUGUUUAGACAAAGUAUGAA | AAG-GUU-GGU  \|\|\|-\|\|\|-:\|\|  UUCUCAAGUCA |
| 75 | GUAUUAAUUCUCAAGUCAAUUGUUUAGACAAAGUAUGAAA |  |
| 76 | UAUUAAUUCUCAAGUCAAUUGUUUAGACAAAGUAUGAAAG | UAAG-GUU-GGU  \|\|\|\|-\|\|\|-:\|\|  AUUCUCAAGUCA |
| 77 | AUUAAUUCUCAAGUCAAUUGUUUAGACAAAGUAUGAAAGU | AG-GUU-GGU  \|\|-\|\|\|-:\|\|  UCUCAAGUCA |
| 78 | GAUAAAUUCAACACGCUCAUAUACUGUGUAACGUCCGUAA | UAAGGUUG  \|\|\|\|-\|\|\|  AUUC-AAC |
| 79 | AUAAAUUCAACACGCUCAUAUACUGUGUAACGUCCGUAAU | AGGUUG  \|\|-\|\|\|  UC-AAC |
| 80 | AAGACGUCAACGAAAAAGUUCUGUGCGGAGGGGUCAGGAA |  |
| 81 | GAUCACUCAACAUUCAUAGUGCGACAGAGUUGCAGAUUAC |  |
| 82 | GUCGUCGUUCCGGUUUGGUGUAAAAAAAGGAGAGACAAUU | AAGG  \|\|\|\|  UUCC |
| 83 | UCGUCGUUCCGGUUUGGUGUAAAAAAAGGAGAGACAAUUC |  |
| 84 | UCUUGUCUUCCCUUUUUCGAUAAUAUUUUGCGUCGUAUCA |  |
| 85 | CUUGUCUUCCCUUUUUCGAUAAUAUUUUGCGUCGUAUCAA |  |
| 86 | AGUAGUAUUCCUUAAAGACGUCAACGAAAAAGUUCUGUGC |  |
| 87 | UACCAUCUUCCUUUAAACUUGGGGACUCUUACAUACUGGA |  |
| 88 | ACCAUCUUCCUUUAAACUUGGGGACUCUUACAUACUGGAU |  |
| 89 | UCGGAUCUUCCGUUUUAAUAAACGGUAUUUGUAAAGGUAG |  |
| 90 | CGGAUCUUCCGUUUUAAUAAACGGUAUUUGUAAAGGUAGU |  |
| 91 | GUGAAAAUUCCGUUCUUGGGAAAAAAUACGUUCUGAUACA |  |
| 92 | UUUACUCCAUUUCGUACCAUGGUUCACGUUACGAAUAGAC | AAGGU-UGGU  \|\|:\|:-\|\|\|\|  UUUCGUACCA |
| 93 | UUACUCCAUUUCGUACCAUGGUUCACGUUACGAAUAGACA |  |
| 94 | ACUCCAUUUCGUACCAUGGUUCACGUUACGAAUAGACACG | UAAGGU-UGGU  \|\|\|:\|:-\|\|\|\|  AUUUCGUACCA |
| 95 | CGUAUCAAUCCUGACGCCUUUCGUAUAAUAUUUUCUUUAA | AGG-UUG  \|\|\|-:\|\|  UCCUGAC |
| 96 | UGUUUACUUCCAAGUACAAUUGAUUGAACGUGUGUUAGUA | AAGGUU  \|\|\|\|\|\|  UUCCAA |
| 97 | GUUUACUUCCAAGUACAAUUGAUUGAACGUGUGUUAGUAG |  |
| 98 | AAUGUAAUUCCAAAAAAUUUUUUAGUUUAUCUGUAGAUUC |  |
| 99 | CUGUAGAUUCCAAGGUUUAGCCUUAAUUUAUAAAAAUUUU |  |
| 100 | AUACAGAUUCCAAAUAAAACAAUCUCGUUGUCGGAUCUUC |  |
| 101 | UAGGUACUUCCAAAGUGAAAAUUCCGUUCUUGGGAAAAAA |  |
| 102 | AGGUACUUCCAAAGUGAAAAUUCCGUUCUUGGGAAAAAAU |  |
| 103 | UUUACUUCCAAGUACAAUUGAUUGAACGUGUGUUAGUAGU | AGGUU  \|\|\|\|\|  UCCAA |
| 104 | GUGACCAUCCAAAAGGUAUAAUGUAAGCCAGAGUGUAACG |  |
| 105 | AAGAAUUCCAAUGUUUAUAUAACUCCGUAAAAAGAGGGUG |  |
| 106 | UGUAAUUCCAAAAAAUUUUUUAGUUUAUCUGUAGAUUCCA |  |
| 107 | GUAGAUUCCAAGGUUUAGCCUUAAUUUAUAAAAAUUUUGU |  |
| 108 | ACAGAUUCCAAAUAAAACAAUCUCGUUGUCGGAUCUUCCG |  |
| 109 | GGUACUUCCAAAGUGAAAAUUCCGUUCUUGGGAAAAAAUA |  |
| 110 | GACGACCUCCAAAUUCGAGGUACCGGAAGAUUGUGAUUCG |  |
| 111 | ACGACCUCCAAAUUCGAGGUACCGGAAGAUUGUGAUUCGU |  |
| 112 | GUAGUAUUCCUUAAAGACGUCAACGAAAAAGUUCUGUGCG | UAAGG  \|\|\|\|\|  AUUCC |
| 113 | UGAAAAUUCCGUUCUUGGGAAAAAAUACGUUCUGAUACAC |  |
| 114 | UGACCAUCCAAAAGGUAUAAUGUAAGCCAGAGUGUAACGU | UAAGGUU  \|\|-\|\|\|\|  AU-CCAA |
| 115 | CGUUAGUGUCCUUGUGUCCCACCGUGUGUCCCGAGAUUUC | AGGUUGG  \|\|\|-\|\|\|  UCCCACC |
| 116 | GUUAGUGUCCUUGUGUCCCACCGUGUGUCCCGAGAUUUCA |  |
| 117 | UAGUGUCCUUGUGUCCCACCGUGUGUCCCGAGAUUUCACC |  |
| 118 | GUCCUUGUGUCCCACCGUGUGUCCCGAGAUUUCACCCCGU |  |
| 119 | UCCUUGUGUCCCACCGUGUGUCCCGAGAUUUCACCCCGUU |  |
| 120 | CCUUGUGUCCCACCGUGUGUCCCGAGAUUUCACCCCGUUG |  |
| 121 | CUUGUGUCCCACCGUGUGUCCCGAGAUUUCACCCCGUUGU |  |
| 122 | CCGAGAUUUCACCCCGUUGUAUCUCACUAAAAAAAAAAGG | UAAGGUUGG  \|\|\|:\|\|-\|\|  AUUUCA-CC |
| 123 | CGAGAUUUCACCCCGUUGUAUCUCACUAAAAAAAAAAGGA | AAGGUUGG  \|\|\|-\|-\|\|  UUC-A-CC |
| 124 | GAUUUCACCCCGUUGUAUCUCACUAAAAAAAAAAGGAAUU | AG-GUUGGU  \|\|-\|\|-\|:\|  UCUCA-CUA |
| 125 | AUUUCACCCCGUUGUAUCUCACUAAAAAAAAAAGGAAUUA |  |
| 126 | GUGUUUUCGCAAGUAUUGGUAAAGUAUACAGAUUGAGCAU | AG-GUU  \|\|-\|\|\|  UCGCAA |
| 127 | UUCACUUUGCCGACUUCAAACGACAACUAUUUUGUAAACU | AA-GGUUG  \|\|-\|\|:\|\|  UUGCCGAC |
| 128 | AAACGACAACUAUUUUGUAAACUUACCAUCUUCCUUUAAA | GUUGGU  \|\|\|\|:\|  CAACUA |
| 129 | GAAGAAUUCCAAUGUUUAUAUAACUCCGUAAAAAGAGGGU | UAAGGUU  \|\|\|\|\|\|\|  AUUCCAA |
| 130 | AUGUAAUUCCAAAAAAUUUUUUAGUUUAUCUGUAGAUUCC |  |
| 131 | UGUAGAUUCCAAGGUUUAGCCUUAAUUUAUAAAAAUUUUG |  |
| 132 | UACAGAUUCCAAAUAAAACAAUCUCGUUGUCGGAUCUUCC |  |
| 133 | ACCGUAGUCCAUCAAAUAUUUAUAAUUCUCUAAACACUAA | AGGUUGGU  \|\|\|\|-:\|\|  UCCA-UCA |
| 134 | CCGUAGUCCAUCAAAUAUUUAUAAUUCUCUAAACACUAAU |  |
| 135 | AUUUAUAAUUCUCUAAACACUAAUUUGAACUAGUGUUUAG | AAG-G-UUGGU  \|\|\|-\|-\|\|-\|\|  UUCUCUAAACA |
| 136 | UUUAUAAUUCUCUAAACACUAAUUUGAACUAGUGUUUAGU |  |
| 137 | UUAUAAUUCUCUAAACACUAAUUUGAACUAGUGUUUAGUG |  |
| 138 | AAUUCAAAUUCUAACCUCUGUUGUUUUUUGUGACCUCUUC | AAGGUUGG  \|\|\|:\|\|\|\|  UUCUAACC |
| 139 | UCAAAUUCUAACCUCUGUUGUUUUUUGUGACCUCUUCGAA | AGGUUGG  \|\|:\|\|\|\|  UCUAACC |
| 140 | CAAAUUCUAACCUCUGUUGUUUUUUGUGACCUCUUCGAAU | GGUUGG  \|:\|\|\|\|  CUAACC |
| 141 | AGUGAGUCCCAAGGAAUCUUUCGUCUCGACUGUCAAAAGA | AGG-UU  \|\|\|-\|\|  UCCCAA |
| 142 | AGAGACACCCACCUACUACGUCUAUUUCGUCCUUCCUGUC | GGUUGG  \|\|\|-\|\|  CCA-CC |
| 143 | UUUCGUCCUUCCUGUCAUAAGUUUACAGUGAAUGUAUAUU | AAGGUUGGU  \|\|\|\|-::\|\|  UUCCUGUCA |
| 144 | UUCGUCCUUCCUGUCAUAAGUUUACAGUGAAUGUAUAUUU |  |
| 145 | UCGUCCUUCCUGUCAUAAGUUUACAGUGAAUGUAUAUUUA |  |
| 146 | UCGUAGUUCAAGUCUUAGUACACGACCCUUUAUACUCUGU | AAGGUU  \|\|\|-\|\|  UUC-AA |
| 147 | GAUUUCUUCAAAAAAAAAAAAAGGAAAAGGUUUCUACCUU |  |
| 148 | UAUUAAUUGCCCCGUUUAACGUCGUUUGGAGAUGACCAUA | UAA-GG  \|\|\|-\|\|  AUUGCC |
| 149 | CCAUAAUAUCAAUGUCACGUUUUUUUUUUUUUAAUUUUAA | UA-AGGUU  \|\|-\|\|-\|\|  AUAUC-AA |
| 150 | UCGAAACCAUCCACUUUAUACGUACGUUGUAAUACUUUUC | GGUUGGU  \|\|\|-\|\|\|  CCAUCCA |
| 151 | AACCAUCCACUUUAUACGUACGUUGUAAUACUUUUCAUAU | GGU  \|\|\|  CCA |
| 152 | GUAAUACUUUUCAUAUCAUUCUUUAAAUACACUCCUCUGA | AAGGU-UGGU  \|\|:\|\|-\|:\|\|  UUUCAUAUCA |
| 153 | AAUACUUUUCAUAUCAUUCUUUAAAUACACUCCUCUGAUU |  |
| 154 | AUACUUUUCAUAUCAUUCUUUAAAUACACUCCUCUGAUUC | AAGGU-UGGU  \|\|\|-\|-\|:\|\|  UUC-AUAUCA |
| 155 | AAAUACACUCCUCUGAUUCCCAUUUCCACGACACAAAACG | AAGGU--UG-GU  \|\|\|\|\|--\|\|-\|\|  UUCCACGACACA |
| 156 | AAUACACUCCUCUGAUUCCCAUUUCCACGACACAAAACGA |  |
| 157 | AUACACUCCUCUGAUUCCCAUUUCCACGACACAAAACGAA |  |
| 158 | UACACUCCUCUGAUUCCCAUUUCCACGACACAAAACGAAG |  |
| 159 | ACACUCCUCUGAUUCCCAUUUCCACGACACAAAACGAAGA |  |
| 160 | UCCUCUGAUUCCCAUUUCCACGACACAAAACGAAGAAUUU |  |
| 161 | CCUCUGAUUCCCAUUUCCACGACACAAAACGAAGAAUUUA |  |
| 162 | CUCUGAUUCCCAUUUCCACGACACAAAACGAAGAAUUUAU |  |
| 163 | UCUGAUUCCCAUUUCCACGACACAAAACGAAGAAUUUAUU |  |
| 164 | CUGAUUCCCAUUUCCACGACACAAAACGAAGAAUUUAUUC |  |
| 165 | UGAUUCCCAUUUCCACGACACAAAACGAAGAAUUUAUUCA |  |
| 166 | GAUUCCCAUUUCCACGACACAAAACGAAGAAUUUAUUCAU |  |
| 167 | AUUCCCAUUUCCACGACACAAAACGAAGAAUUUAUUCAUU |  |
| 168 | UCCCAUUUCCACGACACAAAACGAAGAAUUUAUUCAUUGG |  |
| 169 | UUCCCAUUUCCACGACACAAAACGAAGAAUUUAUUCAUUG | UAA-GGU--UG-GU  \|\|\|-\|\|\|--\|\|-\|\|  AUUUCCACGACACA |
| 170 | CCCAUUUCCACGACACAAAACGAAGAAUUUAUUCAUUGGU | AGGU--UG-GU  \|\|\|\|--\|\|-\|\|  UCCACGACACA |
| 171 | CCAUUUCCACGACACAAAACGAAGAAUUUAUUCAUUGGUU | GGU--UG-GU  \|\|\|--\|\|-\|\|  CCACGACACA |
| 172 | UUAGACGUCGUAUAAAAUCCACACUAUACAGAUUCCAAAU | AGGU-UGGU  \|\|\|\|-\|\|:\|  UCCACACUA |
| 173 | UAGACGUCGUAUAAAAUCCACACUAUACAGAUUCCAAAUA |  |
| 174 | ACAUUAUUCAAUGUUGAUAUGAUCACUCAACAUUCAUAGU | UAAGGUU  \|\|\|\|-\|\|  AUUC-AA |
| 175 | GUCAUAUUGCACCUUUCUUUUCUGUUGUAACCGCGUGAAG | UAAGGUUGG  \|\|\|-\|\|-\|\|  AUUGCA-CC |
| 176 | UUUCGUCUUCACUCCUUUCUUCUAUUCUAACACAAACGCG | AAGGUUG-GU  \|\|\|:\|\|\|-\|\|  UUCUAACACA |
| 177 | UUCGUCUUCACUCCUUUCUUCUAUUCUAACACAAACGCGU |  |
| 178 | UCGUCUUCACUCCUUUCUUCUAUUCUAACACAAACGCGUU |  |
| 179 | CGUCUUCACUCCUUUCUUCUAUUCUAACACAAACGCGUUU |  |
| 180 | GUCUUCACUCCUUUCUUCUAUUCUAACACAAACGCGUUUU |  |
| 181 | UCUUCACUCCUUUCUUCUAUUCUAACACAAACGCGUUUUU |  |
| 182 | CUUCACUCCUUUCUUCUAUUCUAACACAAACGCGUUUUUU |  |
| 183 | UUCACUCCUUUCUUCUAUUCUAACACAAACGCGUUUUUUG |  |
| 184 | UCACUCCUUUCUUCUAUUCUAACACAAACGCGUUUUUUGA |  |
| 185 | ACUCCUUUCUUCUAUUCUAACACAAACGCGUUUUUUGAUU |  |
| 186 | UCCUUUCUUCUAUUCUAACACAAACGCGUUUUUUGAUUUU |  |
| 187 | UUUCUUCUAUUCUAACACAAACGCGUUUUUUGAUUUUGAU |  |
| 188 | UUCUUCUAUUCUAACACAAACGCGUUUUUUGAUUUUGAUG |  |
| 189 | UCUUCUAUUCUAACACAAACGCGUUUUUUGAUUUUGAUGU |  |
| 190 | CUUCUAUUCUAACACAAACGCGUUUUUUGAUUUUGAUGUU |  |
| 191 | UCUAUUCUAACACAAACGCGUUUUUUGAUUUUGAUGUUAA | GGUUG-GU  \|:\|\|\|-\|\|  CUAACACA |

Mimosa identified 12 canonical functional segments (highlighted in blue), within which we found a total of 7 non-overlapping target sites, including:

**Site A:**

UAGGUAUUCCACGUCUGACGUUUAAAGACGUCGUACUAAU

**Site B:**

UGUUUACUUCCAAGUACAAUUGAUUGAACGUGUGUUAGUA

GUUUACUUCCAAGUACAAUUGAUUGAACGUGUGUUAGUAG

**Site C**:

GAAGAAUUCCAAUGUUUAUAUAACUCCGUAAAAAGAGGGU

**Site D:**

AAUGUAAUUCCAAAAAAUUUUUUAGUUUAUCUGUAGAUUC

AUGUAAUUCCAAAAAAUUUUUUAGUUUAUCUGUAGAUUCC

**Site E:**

CUGUAGAUUCCAAGGUUUAGCCUUAAUUUAUAAAAAUUUU UGUAGAUUCCAAGGUUUAGCCUUAAUUUAUAAAAAUUUUG

**Site F:**

AUACAGAUUCCAAAUAAAACAAUCUCGUUGUCGGAUCUUC UACAGAUUCCAAAUAAAACAAUCUCGUUGUCGGAUCUUCC

Site G:

UAGGUACUUCCAAAGUGAAAAUUCCGUUCUUGGGAAAAAA AGGUACUUCCAAAGUGAAAAUUCCGUUCUUGGGAAAAAAU
